# ProbeST: a custom probe design pipeline for Spatial Transcriptomics

**DOI:** 10.1101/2025.08.29.673037

**Authors:** Sofia Rouot, Ireen van Dolderen, Patrick Rosendahl Andreassen, Solène Frapard, Sybil A. Herrera-Foessel, Hailey Sounart, Sami Saarenpää, Julia A. Vorholt, Stefania Giacomello

**Author notes:** Corresponding author: Stefania Giacomello. First-shared author (Sofia Rouot, Ireen van Dolderen, Patrick Rosendahl Andreassen). www.re-peat.earth.

## Abstract

Probe-based Spatial Transcriptomics profiles spatially-resolved transcriptomes using gene-specific probe pairs for both Formalin-fixed paraffin-embedded (FFPE) and Fresh Frozen samples. However, its applicability is restricted to human and mouse studies, due to commercial probe set availability. Here, we present ProbeST, an open-source computational pipeline for designing custom probe sets for any organism of interest. We validated ProbeST on FFPE mouse enteroids infected with *Salmonella enterica* serovar Typhimurium, using custom pathogen probes with the available mouse probe panel. We simultaneously detected host and pathogen transcripts, demonstrating high probe specificity and enabling identification of inflammatory response host genes *Mefv*, *Tnf*, and *Anxa1* colocalizing to pathogen genes. The reproducible ProbeST workflow expands probe-based Spatial Transcriptomics to studies of non-model organisms and host-pathogen interactions.

## Background

The Spatial Transcriptomics technology (ST; 10X Genomics Visium) combines high-resolution histological imaging with sequencing-based 2D gene expression maps (1). The original poly(A) capture approach (1,2), compatible with Fresh Frozen (FF) tissue samples, underperforms in clinically-relevant FFPE tissue due to formalin-induced mRNA degradation and cross-linking between RNA and other biomolecules (3,4). To address these challenges, the Visium FFPE-compatible approach maintains sensitive transcript detection through gene-specific probe pairs (5), which consist of a Left-Hand Side (LHS) and a Right-Hand Side (RHS), each containing a hybridizing sequence and a probe handle (**Fig. 1.A)**. The LHS and RHS of a probe pair ligate only if they bind adjacent to each other on the target mRNA. After RNA degradation, the ligated probe pair is captured onto the Visium slide via the poly(A) tail on the RHS probe handle. This probe-based chemistry, despite requiring prior knowledge of gene sequences to design probe pairs, enables spatial profiling of whole transcriptomes and non-polyadenylated transcripts. Currently, commercially available probe sets exist only for the human and mouse whole transcriptomes, covering ∼18,000 and ∼19,000 genes, respectively. Therefore, custom probe design is essential to extend Visium FFPE to the study of additional organisms (6,7).

**Figure 1:**
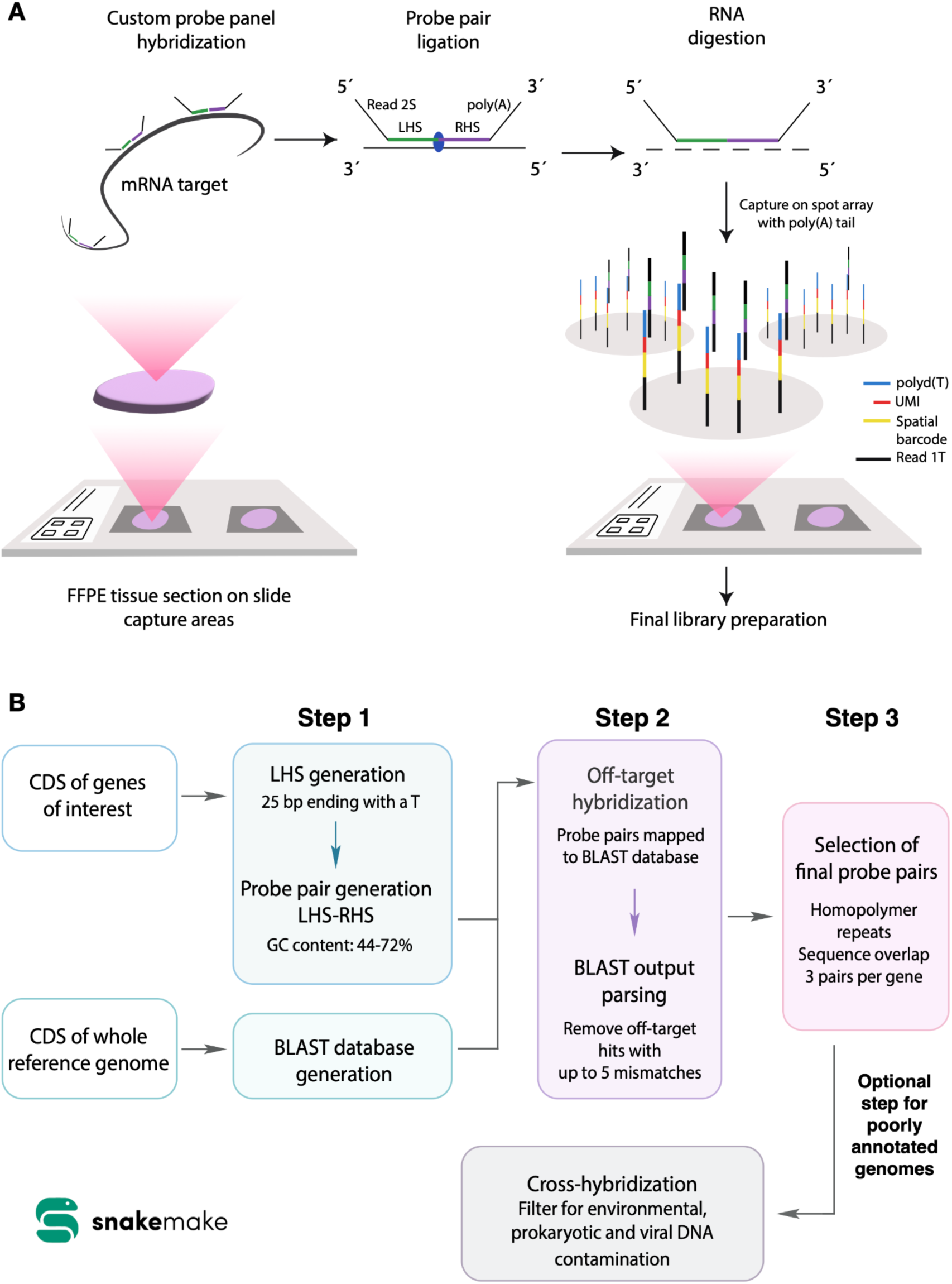
An overview of the probe-based chemistry and the user-friendly ProbeST pipeline. A) Main steps of the ST probe-based chemistry with a custom probe panel. For each mRNA target, up to three probe pairs are custom designed. Each probe consists of a 25 bp sequence reverse complementary to the sequence of the targeted transcript. The probe handle on the Left-Hand Side (LHS) probe contains the partial read 2S sequence used for downstream sequencing. The probe handle on the Right-Hand Side (RHS) probe consists of a poly(A) tail for probe capture on the Visium spot array. After probe hybridization, the probe sides are ligated to form a 50 bp probe pair. The mRNA undergoes digestion, and the probe pairs are captured by the spatial probes at specific spots of the array. Subsequent steps are performed to obtain final libraries ready for sequencing. Custom probes can be designed for any organism of interest. **B) General workflow of the ProbeST pipeline.** Two input FASTA files are required. The first input file is used to generate potential probe pairs, while the second input file is used to create a BLAST database. The generated potential probe pairs are then assessed for off-target hybridization against the BLAST database. The probe pairs that match to more than two sites in the genome with fewer than six mismatches are removed. If there are more than six mismatches, the match is considered insufficiently specific, resulting in a low likelihood of the probe pair binding to this target. The resulting output is parsed and up to three non-overlapping probe pairs per gene are selected for high specificity. An optional add-on cross-hybridisation step for poorly annotated genomes removes probes with over 99% identity against the NCBI’s environmental, prokaryotic and viral databases.

We present ProbeST, an open-source and scalable pipeline to custom design probe pairs compatible with the probe-based Visium assays (8). We experimentally validated ProbeST by designing probe pairs specific to coding regions of the *Salmonella enterica* subsp. *enterica* serovar Typhimurium strain SL1344 (*S*. Tm) and combined them with the mouse whole transcriptome probe set to analyze adult derived mouse intestinal organoids (enteroids) infected by *S*. Tm with high specificity, thus enabling the study of specific host-pathogen interactions. ProbeST is publicly available on GitHub as an automated SnakeMake workflow, allowing reproducible and ease-of-use custom probe design for ST studies beyond human and mouse to any eukaryotic and prokaryotic organism (9).

## Results and Discussion

### ProbeST custom probe design pipeline

We developed ProbeST to address the need for an automated custom probe design pipeline for Visium FFPE studies of any organism of interest. ProbeST is designed to output a final probe set optimized for specific hybridization to target genes by meeting the following guidelines (as suggested by 10X Genomics probe design instructions (8)): the probes should target coding regions, the 25th nucleotide of the LHS probe (at the 3’ end) must be a T, and the GC content should be between 44-72% for each probe (**Fig. 1.B)**.

ProbeST consists of three main steps integrated into a user-friendly SnakeMake pipeline. In step 1, the pipeline first utilizes Primer3, an open-source program for designing hybridization probes (10), to generate 25 bp LHS probe sequences complementary to the Coding DNA Sequences (CDS) in the first input file, and then extends these sequences into 50 bp probe pairs (LHS + RHS). In step 2, ProbeST uses the second input file to include in a BLAST step that removes probe pairs with off-target hybridization on the reference transcriptome (11), corresponding to probe pairs with at least two BLAST hits of less than six mismatches (8). In step 3, the pipeline selects the final probe pair set by retaining up to three non-overlapping probe pairs per gene, while excluding probes with homopolymeric regions longer than 5 bp (**Fig. 1.B)**. ProbeST generates three files as final outputs, including a file formatted for use as input for the Space Ranger software, the bioinformatics tool used to analyze and process Visium data (12).

An optional add-on step can be included to remove sequences in the probe pairs that arise from cross-hybridization, defined as contamination from the environment. Indeed, poorly annotated genomes often contain sequences from environmental, viral, or prokaryotic DNA (13,14). The cross-hybridization step keeps probe pairs of higher specificity to the organism of interest and filters them based on the same criteria as the general ProbeST pipeline (8). Overall, ProbeST designs highly specific custom probe pairs that exclusively target the gene transcripts, including non-polyadenylated transcripts.

### ProbeST enables the Dual Spatial Transcriptomics study of *S*. Tm infection in mouse enteroids

To assess the ProbeST pipeline, we applied it to study the *Salmonella enterica* subsp. *enterica* serovar Typhimurium strain SL1344 (*S*. Tm) infection process and the intestinal host response to pathogen invasion leveraging mouse intestinal organoids (enteroids). We selected 19 genes from a previous study that investigated the transcriptional adaptations of *Salmonella enterica* during infection (15). We used the *S*. Tm CDS of the selected genes as input for ProbeST, with the header format modified (see Methods), and generated a total of 51 *S*. Tm probe pairs (**Additional files 1-3, Table 1**).

The gene panel included markers of *S*. Tm’s cytosolic and vacuolar life cycles, representing different stages of *S*. Tm infection (**Table 2**). Indeed, during *S*. Tm infection of the small intestine, the pathogen invades intestinal epithelial cells (IECs), adopts a vacuolar phase, and, depending on host cell conditions, can potentially transition to a cytosolic phase. The host IEC layer senses the invasion through the NAIP/NLRC4 inflammasome, leading to Caspase-1 activation, Gasdermin D-mediated pore formation, and cytokine IL-18 release (16). The panel also included *S*. Tm housekeeping genes to capture highly abundant transcripts, as *S*. Tm transcripts are less abundant than host transcripts in the tissue.

To study the *S*. Tm infection process in a mouse model and validate our custom probes, we applied dual Spatial Transcriptomics (DualST), an approach that uses two probe sets to simultaneously detect spatially resolved transcripts from both host and pathogen (6). Specifically, we targeted 19,405 mouse genes with the commercially available 55,538 mouse probes and 19 *S*. Tm genes with our custom-designed 51 *S*. Tm probes, corresponding to a 1089-fold excess of host probes. We applied DualST to four monolayer mouse enteroid samples, representing different experimental conditions: wild-type non-infected (WT), WT infected (WT+), *Nlrc4*^-/-^ infected, and *GsdmD*^-/-^ infected, with an *S*. Tm infection load equivalent to about 500 bacteria across all samples. *Nlrc4* is an inflammasome sensor that detects intracellular bacterial components, triggering epithelial defense responses including interleukin maturation, cell expulsion from the monolayer and GSDMD cleavage. Cleaved GSDMD N-terminal forms pores in the membrane that allows mature interleukins to be secreted and thus contributes to pyroptotic cell death by amplifying host inflammation (17). These samples were designed to assess the effect of *S*. Tm infection on host transcriptional responses and the contribution of *Nlrc4* and *GsdmD* for those.

First, we tested the specificity of the ProbeST probe set to detect *S*. Tm transcripts by comparing *S*. Tm signals across infected conditions (WT+, *Nlrc4^-/-^*, *GsdmD^-/-^*) and the control condition (WT). We observed a high number of *S*. Tm UMIs in the infected conditions (482 UMIs for WT+, 562 for *GsdmD^-/-^* and 766 for *Nlrc4^-/-^*), compared to only 24 UMIs in WT (Fig 2**.A)**. The low background signal observed in the latter can be explained by unspecific binding. We then verified probe sensitivity through the dual capture of host and low-abundance *S*. Tm transcripts in the infected samples (Fig. 2**.B)**. The detection of *S*. Tm transcripts demonstrates that the ProbeST custom probes hybridize to the target genes, undergo ligation, and are detected with host probes, despite the 1089-fold difference in probe abundance. We observed that the total UMI counts per *S*. Tm gene varied considerably, from 764 UMIs for *sipC* and 340 for *lpp,* to 1 UMI for *uhpT* and *zinT* (Fig. 2**.C)**. The UMI variation within this gene set has a strong Spearman correlation (0.683) with the Transcript per Million (TPM) observed in an earlier study (15) **(Additional file 4)**, and may reflect differences in transcript abundance.

**Figure 2:**
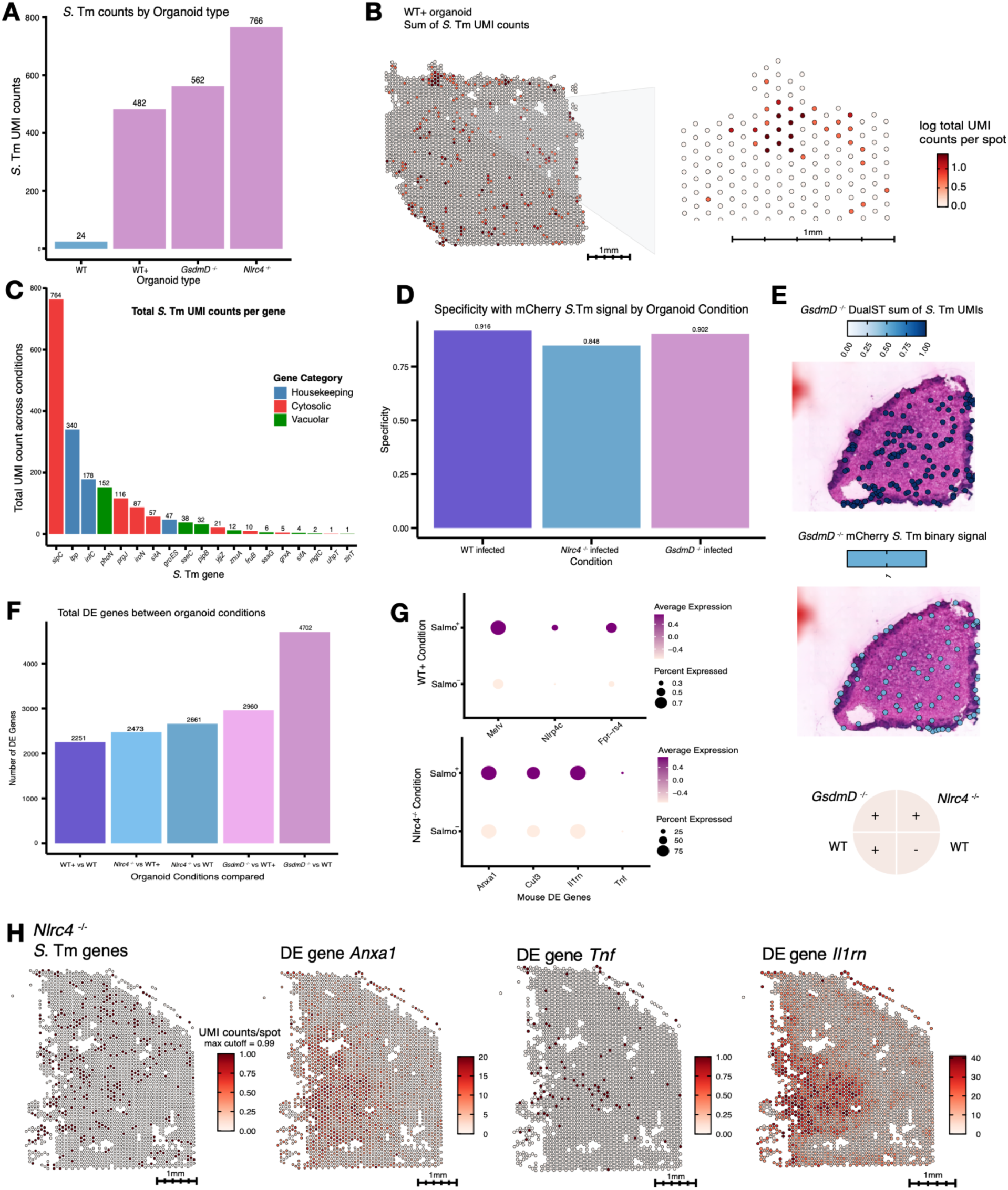
Application of ProbeST for DualST data of *S*. Tm infection in mouse enteroids. **A)** *S*. Tm detection by the ProbeST custom probe panel in the three infected enteroid conditions: wild-type infected (WT+), *Nlrc4*^-/-^ infected, and *GsdmD*^-/-^ infected (pink), compared to the control non-infected enteroid WT (blue). The sum of UMI counts across all *S*. Tm unique gene transcripts was calculated for each condition. **B)** Visualization of *S*. Tm detection spatially mapped onto the sample WT+. The UMI counts represent the total UMIs per spot of all the captured *S*. Tm unique gene transcripts. The values shown are log1p-normalized with parameter max_cutoff = 0.99. **C)** Plot of the total UMI counts for each *S*. Tm gene across all spots in all 4 conditions. **D)** Validation of the *S*. Tm DualST signal with the fluorescent mCherry signal. The mCherry spatial coordinates were aligned to the DualST coordinates with a computational approach. Specificity was calculated for each infected condition through a confusion matrix, with the mCherry signal considered as ground-truth. **E)** The custom probe *S*. Tm DualST coordinates (top) and the mCherry coordinates (middle) mapped onto the *GsdmD*^-/-^ infected condition, with the four-sample layout (bottom). **F)** Total number of mouse DE genes between different enteroid conditions. **G)** Colocalization analysis. Dot plot of the mouse DE genes between spots with *S*. Tm probe capture (Salmo^+^) and spots without *S*. Tm capture (Salmo^-^) in the WT+ and *Nlrc4^-/-^* conditions. **H)** Colocalization analysis. *S*. Tm genes and selected mouse DE genes spatially mapped onto the *Nlrc4^-/-^* condition. UMI = Unique Molecular Identifier; DE = Differentially Expressed.

Next, we validated *S*. Tm custom probe detection by comparing the DualST signal with mCherry fluorescence, a constitutively expressed red fluorescent protein in the *S*. Tm used for infection (17). We developed a computational approach to compare spot coordinates from DualST data with pixel coordinates from fluorescence imaging (see Methods). Across all three enteroid conditions, we observed a spatial overlap with high specificity (WT+: 0.916, *GsdmD*^-/-^: 0.848, and *Nlrc4*^-/-^: 0.902) and lower sensitivity (WT+: 0.167, *GsdmD*^-/-^: 0.163, and *Nlrc4*^-/-^: 0,117) (Fig. 2**.D-E)**. The high specificity confirmed that the spatial distributions of the DualST and the mCherry signals match. The lower sensitivity likely resulted from the different resolutions, as mCherry fluorescence is not at single-molecule resolution and is inherently less sensitive than probe-based detection. Moreover, a likely explanation for the low sensitivity is potential competition of probe sequences on the target mRNA. Indeed, we found that for genes with more than 1 probe pair, 0-90% of the 50 bp sequences overlapped **(Additional file 5)**. We addressed this issue in the ProbeST pipeline, reducing the overlap to 0% between probe pairs targeting the same gene. The updated version is also available on GitHub. Overall, the spatial overlap between fluorescent and probe *S*. Tm signals validated that custom probes captured *S*. Tm transcript distribution with high specificity.

### Host transcriptional responses are condition-specific during *S*. Tm infection

To investigate enteroid responses to *S*. Tm infection, we performed Differential Expression Analysis (DEA) between WT and each of the infected enteroid conditions (WT+, *GsdmD^-/-^*, and *Nlrc4^-/-^*). The comparison between infected *GsdmD^-/-^* and uninfected WT enteroids yielded 4702 significantly differentially expressed (DE) genes (adjusted p-value < 0.05), the highest among all comparisons, followed by 2960 DE genes observed between infected *GsdmD^-/-^* and infected WT+ enteroids (Fig. 2**.F, Additional files 6-10)**. While DE gene counts do not directly reflect infection severity, these differences suggest distinct transcriptional responses in the absence of *GsdmD* and *Nlrc4*. *GsdmD* encodes a pore-forming effector involved in pyroptotic cell death downstream of inflammasome activation, and its deletion may alter IEC death and inflammatory signaling dynamics (18). NLRC4, in contrast, acts upstream as a pattern recognition receptor sensing bacterial flagellin and type III secretion system components, triggering inflammasome activation and downstream responses, including cytokine release and, in IECs, epithelial cell extrusion.

The altered gene expression profiles observed in *GsdmD^-/-^* and *Nlrc4^-/-^* enteroids following *S*. Tm infection likely reflect the consequences of knocking-out these genes, which can be captured with this method. These transcriptional changes may suggest potential effects on epithelial integrity and pathogen control, although confirming such implications would require further functional insight.

While only one biological sample per condition was included, the spatial resolution of the data allows for an exploratory comparison of transcriptional trends across conditions. We observed significant downregulation of the transcription factors *Stat3*, *Fos*, *Fosl1*, *Egr1*, and *Atf3*, in both *GsdmD^-/-^* and *Nlrc4^-/-^* enteroids compared to WT and WT+ enteroids. These transcription factors are known to be induced during early *S*. Tm infection and to regulate inflammatory and stress response pathways in epithelial cells (17,19,20). Consistent with these findings, the GO term “STAT Family Protein Binding” (GO:0097677) showed significant downregulation in the *Nlrc4^-/-^* and WT+ comparison **(Additional file 11)**, suggesting attenuation of STAT-mediated signaling in the absence of NLRC4.

In contrast, the transcriptional differences between WT+ and WT enteroids were relatively modest. A likely explanation is the immature state of IECs in monolayer culture, which has been reported to dampen inflammasome activity following NLRC4 activation (21). Nevertheless, several biological processes previously associated with host epithelial defense including regulation of nitric oxide metabolism, lysozyme production, and peroxisomal transport were upregulated in WT+ enteroid relative to WT enteroid **(Additional file 12)** (22–25).

Taken together, these spatially resolved transcriptional patterns are consistent with an altered epithelial response to *S.* Tm infection in the absence of NLRC4 or GSDMD. While biological replication is needed to confirm these findings, the observed differences suggest that inflammasome signaling contributes to shaping the early transcriptional landscape of infected enteroids and demonstrates that DE genes can be detected between conditions. To further investigate host-pathogen interactions at the spatial level, we performed a colocalization analysis to examine localized host transcriptional changes at *S.* Tm infection sites, enabled by our ProbeST custom probe set.

### Host-pathogen colocalization analysis highlights the role of the innate immune system in *S*. Tm-infected IECs

To study the spatial and local effect of *S*. Tm infection on the mouse IEC transcriptional profile, we performed a colocalization analysis comparing gene expression profiles between spots that contained *S.* Tm signal (Salmo^+^, at least 1 UMI of any *S*. Tm gene) and spots with zero *S.* Tm signal (Salmo^-^).

In the WT+ condition, DEA and pathway analysis (26) revealed upregulation of *Mefv*, *Nlrp4c*, and *Fpr-rs4* (adjusted p-value < 0.05), genes involved in inflammatory responses (Fig. 2**.G, Additional file 13)**. *Mefv* encodes pyrin, a key regulator of caspase-1 and IL-18 expression in innate immunity (27,28), while *Nlrp4c* is a gene of unknown function that is part of the Nlrp4 family (29), some members of which have been suggested to be part of autoimmune responses.

In the *Nlrc4^-/-^* condition, the genes *Anxa1*, *Cul3*, *Il1rn*, and *Tnf* were upregulated in Salmo^+^ spots (Fig. 2**.G-H, Additional file 14)**. Notably, *Anxa1* plays a regulatory role during intestinal mucosal inflammation (30), and *Tnf* is known to be induced by *S*. Tm flagellin upon intestinal invasion, acting as a proinflammatory signaling molecule for neighboring cells (31). These findings suggest that inflammatory response genes are consistently activated in response to *S.* Tm infection in both WT+ and *Nlrc4^-/-^*Salmo^+^ spots. Finally, the *GsdmD^-/^*^-^ Salmo^+^ spots revealed upregulation of *Tunar*, *Gsd3*, *Aadacl4*, and *Hsf2*, genes with predictive transmembrane helix domains **(Additional file 15)**. Given the involvement of *GsdmD* in forming pores in the infected IEC membrane, the four genes *Tunar*, *Gsd3*, *Aadacl4*, and *Hsf2* might be upregulated as a result of the lack of *GsdmD* expression or pore-formation. Together, these differential gene expressions in innate immune regulators highlight condition-specific local host responses to *S. Tm* infection.

Overall, we present ProbeST, a SnakeMake pipeline that automates and standardizes the design of custom gene-specific probe sets for ST studies of any organism in a reproducible manner. ProbeST minimizes manual input and enables the spatial capture of both polyadenylated and non-polyadenylated transcripts, supporting studies of non-model organisms and host-pathogen interactions. While the current ProbeST version does not yet account for parameters such as common SNPs or melting temperature (Tm) differences between on-target and off-target hybridization, future updates aim to improve design precision by incorporating such features.

Alternative pipelines have been developed for imaging-based ST, including the pipeline Spapros that covers both gene set selection and probe design (32). Though similar in structure, ProbeST is tailored for Visium, the most widely used spatially-resolved transcriptomics technology (33), and enables multi-organism transcript detection. ProbeST’s flexible workflow allows for potential adaptation to other ST platforms, including imaging-based methods, by modifying parameters such as handle sequence and probe length.

We demonstrate the utility of custom probes generated by ProbeST in a DualST experiment, where we detect non-polyadenylated transcripts of the bacteria *S*. Tm in infected mouse enteroids, despite a lower number of prokaryotic transcripts and probes compared to the host. The DualST results corroborate the key role of the innate immunity in the host transcriptional responses to *S*. Tm infection. Our study validates the high specificity and effectiveness of the ProbeST custom probes, demonstrating the pipeline’s potential to detect transcripts from multiple organisms in a single tissue section, and to advance ST probe-based studies beyond mouse and human organisms.

## Conclusions

ProbeST is a custom design SnakeMake pipeline for generating specific probe panels for the probe-based Visium chemistry that can be applied on both FF and FFPE tissue sections, aimed at maintaining spatial transcript retrieval. ProbeST facilitates performing Visium experiments with gene-specific binding probes for organisms other than human or mouse, making it highly applicable for the study of other eukaryotes, microorganisms, and any non-model organism.

The DualST dataset generated to study host-pathogen interactions during *S*. Tm infection of a mouse enteroid validated the specificity of the binding probes by targeting specific gene transcripts of interest. ProbeST, though, has the potential to achieve scalability to the whole transcriptome of any organism, providing a novel way of applying Visium FFPE experiments to study spatial transcriptomics across kingdoms.

## Methods

### ProbeST

#### Input files

ProbeST takes as input the Coding DNA Sequence (CDS) in FASTA file format, trimmed for the genes of interest. The input file consists of a header and its corresponding sequence for each gene transcript, with each header manually modified to contain at least the gene ID, the gene name and the transcript ID in the following order and format:

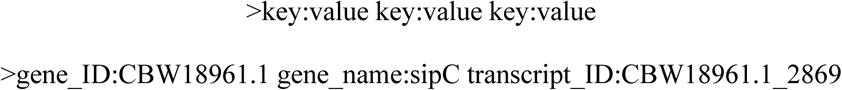

As the CDS file formatting may vary between databases and species, manual editing of the file prior to running ProbeST is necessary. The CDS headers of the 19 *S*. Tm selected genes were modified accordingly and used for the ProbeST pipeline. In total, 51 probe pairs were obtained.

#### Cross-hybridization

The optional cross-hybridization pipeline takes as input the probe pairs from the ProbeST pipeline. The 50 bp probe pair binding sequences are then BLASTed against the environmental, prokaryotic and virus databases downloaded from NCBI. The hits with over 99% similarity are excluded from the final probe pair selection.

### DualST

#### Sample generation

##### Enteroid culture and 2D monolayer preparation

Enteroids were derived from adult C57BL/6 mice isolated in previous studies (21). Cryo-stocked enteroids were revived by thawing in a water bath at 37°C and centrifuged in 5 mL DMEM/F-12 at 200 RCF for 5 minutes. The supernatant was discarded, and the crypts were resuspended in Matrigel (34). Approximately 50 µL of the Matrigel-crypt suspension was plated per well in pre-warmed 24-well plates, followed by incubation at 37°C for 10 minutes to solidify the Matrigel. The embedded crypts were overlaid with 700 µL of Mouse IntestiCult™ Organoid Growth Medium and cultured at 37°C in a 5% CO₂ incubator. The medium was replaced every 2-3 days and passaged every 5-7 days as described previously (35). At least 2 passages were done before revived enteroids were used for sample preparations as described below.

For the preparation of the enteroid 2D monolayer, enteroids were first enriched for proliferative cells by growing them in Human IntestiCult™ Organoid Growth Medium for 4 days after passage. To prepare the growth surface, an Ibidi 4-well removable inserts was placed on a microscope slide. The area in the wells were then coated with 5 µL/cm² Matrigel and allowed to solidify at 37°C for 1 hour. Enteroids enriched for proliferative cells were harvested in ice-cold DMEM/F-12 and spun at 200 rcf to remove the Matrigel and dissociated into single cells by resuspending and incubating in TrypLE Express for 5 minutes at 37°C, followed by gentle pipetting. Dissociated cells were centrifuged at 300 rcf for 5 minutes, and the pellet was resuspended in 100 µL of Human IntestiCult™ Organoid Growth Medium supplemented with 10 µM Y-27632. Approximately 10⁵ cells were seeded into each well of the ibidi 4-well removable insert, on top of the Matrigel-coated microscope slides. Cells were incubated at 37°C in a 5% CO₂ incubator for 4 days with a medium exchange at day 3. Monolayers were monitored under a microscope to ensure confluency and cell health.

##### Bacterial Infection and FFPE preservation

The enteroid monolayers were washed with DMEM/F-12 medium. *Salmonella enterica* serovar Typhimurium strain SL1344/pFPV25.5 (constitutive mCherry) was grown overnight in LB medium (36). The bacterial culture was then diluted 1:100 in fresh LB medium and incubated on a rolling wheel for 4 hours at 37°C. Optical density (OD) was measured to determine bacterial concentration, and 500 or 50 bacteria in 100 µL of Human IntestiCult™ Organoid Growth Medium (verified by plating on LB agar plates) were used to infect each cell layer. The infection was carried out for 2 hours at 37°C. Following infection, the medium was removed, and samples were fixed in 4% paraformaldehyde (PFA). The samples were imaged (see Visual detection section below) and dehydrated by adding a series of 70%, 95% and 100% ethanol for 5 min. Dehydrated samples were embedded in paraffin by heating the microscopy slide to 90°C, adding 100 µL molten paraffin and placing a 90°C clean microscopy slide on top of the paraffin to spread the paraffin thin. The top slide was removed and the sample slide with a thin layer of paraffin was cooled down and solidified. Samples were stored at −80°C until used.

##### Visual detection of bacteria in infected monolayers

Imaging of the infected monolayers was performed on fixed samples in PBS with 1 U/mL RNase inhibitor (Takara) using a Zeiss Celldiscoverer 7 platform and imaged with a PlanApochromat 5x/0.35/WD: 5.1 mm objective captured on an Axiocam 712 mono camera. Entire monolayers were imaged in both brightfield and red fluorescence channels. Red fluorescence was excited using Calibri.2 LED Illumination Diodelaser 561 nm 10 mW, and emission was filtered with a 515/30 nm bandpass filter. The imaging was conducted across 7 focal planes to capture the full depth of the monolayers. Image analysis was performed in FIJI (37). Potential *S.* Tm sites were automatically assigned by subtracting a Z-projection of the median intensity from a Z-projection of max intensity, followed by particle analysis filtering for circularity and size. Automatically assigned *S.* Tm sites were manually curated in the original images containing all 7 focal planes and filtered for actual *S.* Tm sites.

##### Spatial Transcriptomics

The experimental workflow consisted of the CytAssist-enabled FFPE Visium, optimized for the mouse enteroid [18]. The main steps in the workflow are: tissue preparation and imaging, probe mix addition, CytAssist-enabled RNA digestion, probe release, reverse transcription, library preparation and Illumina sequencing (38,39).

Prior to the probe hybridization step 1.1 of the User Guide (39), the *S*. Tm custom probes, ordered as oPool Oligos (50 pmol/oligo scale), were first resuspended in 62.5 µL IDTE (pH 8.0) to a concentration of 800 nM/oligo. A spike-in pool working stock was then prepared with 3 µL of each of the LHS and RHS resuspended stocks and completed to 100 µL with nuclease-free water for a final concentration of 24 nM/probe. The probe mix was prepared with 10 µL of the working stock, 10 µL of RHS Mouse Probes, 10 µL of LHS Mouse Probes, and 70 µL of FFPE Hybridization buffer (8).

##### Spatial Transcriptomics data generation

The gene expression count matrices were generated by space ranger ‘count’ (version 3.0.0) (12). A custom reference file was made from space ranger ‘mkref’ using the Mouse reference dataset (GRCm38 Reference - 2020-A) and the *S*. Tm CDS file (GenBank Assembly GCA000210855.2). A custom probe set was used, by appending the *S*. Tm custom probes to the Mouse Probe Set from 10X Genomics (Visium Mouse Transcriptome Probe Set v1.0).

From this step, data processing and analysis were performed in R (version 4.3.2). The package Semla (version 1.1.6) was used for its tools developed specifically for Spatially Resolved Transcriptomics data analysis and visualization (40).

##### Validation of the S. Tm probes with mCherry fluorescence

The *S.* Tm spatial distribution between the DualST data and the mCherry signal were compared. Firstly, brightfield images were acquired prior to the ST workflow to obtain visual data on *S.* Tm localization in the mouse enteroid monolayers. Following imaging, potential *S.* Tm locations were automatically detected based on mCherry signal intensity and size. Each site was then manually assessed to remove false positives. Pixel coordinates with mCherry red signal were recorded, and the images were vertically flipped to match the DualST images.

Then, the brightfield mCherry images were aligned to the DualST image with the Loupe Browser tool v7.0.1 (41), and the pixel coordinates with mCherry signal were transformed with the resulting transformation matrix. The fluorescent and DualST data come from the same samples, so it was not necessary to adjust for the use of consecutive sections. After alignment and transformation, the pixel coordinates were read as a Seurat object and subsequently merged with the DualST Seurat object. Each pixel coordinate was attributed to the closest Visium spot coordinate within a radius of 100 µm, based on the k-nearest neighbors approach. The DualST data was split into Salmo^+^ and Salmo^-^ spots, with Salmo^+^ corresponding to spots with at least 1 UMI from any *S*. Tm gene. The coordinates of the DualST Salmo^+^ spots were compared with the mCherry spot coordinates.

A confusion matrix was generated, taking the mCherry data as reference. The specificity of the DualST to detect *S*. Tm was calculated with the following formula:

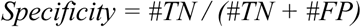

With #TN the spot coordinates with neither mCherry nor DualST signal, and #FP the number of spots with DualST signal but no fluorescent signal.

##### DEA between enteroid conditions and Colocalization analysis

The mouse gene expression data was first filtered and normalized with the Seurat (version 5.0.2) module SCTransform v2 (42), then Differential Expression Analysis (DEA) was performed on mouse genes with the Seurat ‘FindMarkers’ function and the Wilcoxon rank-sum test. More specifically, the conditions WT+, *GsdmD^-/^*^-^, *Nlrc4^-/-^* were each compared to WT to analyze the different effects of *S*. Tm infection in relation to the non-infected control. Then, *GsdmD^-/-^* and *Nlrc4^-/-^* were each compared to WT+ to analyze the effects of the gene knockouts on the enteroid defense response to *S*. Tm infection.

The GO terms for each list of DE genes were obtained with the EnrichR package for the libraries “GO_Biological_Processes_2023”, “KEGG_2019_Mouse”, and “GO_Molecular_Functions_2023” (version 3.4) (43).

The colocalization analysis was performed with a DEA between Salmo^+^ and Salmo^-^ spots, for each enteroid condition WT, WT+, *GsdmD^-/-^*, and *Nlrc4^-/-^*, where Salmo^+^ is attributed to spots with at least 1 UMI from any *S*. Tm gene. The same parameters as above were applied for the DEA. Pathway analysis was performed with the tool DAVID functional annotation tool (v2023q4) (26).

## Declarations

### Ethics approval and consent to participate

No mice were sacrificed for this publication, as all enteroids are derived from another study (18,21,44).

### Consent for publication

Not applicable

### Availability of data and materials

ProbeST is implemented as a Snakemake pipeline and is available on our GitHub repository with its related scripts (https://github.com/giacomellolab/ProbeST) (9). The scripts for the data analysis can be accessed on our GitHub repository.

Sequencing data have been deposited at NCBI-SRA under the BioProject PRJNA1218431. The gene count matrices as well as the high resolution ST H&E images are available on Figshare (45,46).

## Authors’ contributions

S.G., P.R.A. and J.A.V. were responsible for the conceptualization of the study, and secured the necessary funding for the project. S.R., I.v.D., and S.H.F. developed the ProbeST computational pipeline, and S.S. and H.S. helped in its design. P.R.A, S.F., and S.R. conducted the research and investigation process. P.R.A. designed and provided the mouse intestinal organoid samples, collected and analyzed the mCherry-fluorescent imaging data of the pathogen *S*. Tm. S.R. designed the custom probes for *S*. Tm. S.F. and S.R. conducted the CytAssist-enabled DualST experiments. S.R. curated the DualST sequencing data and performed the formal analysis of the data. S.R. and P.R.A. validated the results. S.R. created the visualizations and wrote the original draft of the manuscript. S.G., H.S., S.S., P.R.A. and J.A.V. contributed to the review and editing of the manuscript. S.F. and P.R.A. carried out the project administration and responsibility for the research planning and execution. Study supervision was carried out by S.G.

## Acknowledgments

We would like to thank the Scientific Center for Optical and Electron Microscopy (ScopeM) of ETH Zurich for providing helpful technical advice. The authors thank Michael Stebler (ScopeM, Department of Health Sciences and Technology, ETH Zurich, Switzerland) for his support and assistance with the ZEISS Celldiscoverer 7, Lovisa Franzén for her guidance in the data analysis, and Jeremie Charbord for helpful feedback on the manuscript.

## Figures, tables and additional files

Table 1: Complete list of probe pairs with their corresponding 50 bp sequence.

Table 2: Genes of interest of *S*. Tm targeted for the custom probe design.

Additional file 1: probe_set.csv

Additional file 2: selected_probes.txt

Additional file 3: probe_quantifications.txt

Additional file 4: STm_RNA_seq_TPM_to_DualST_UMIs.xlsx

Additional file 5: Probe_sequences_overlap_summary.xlsx

Additional file 6: DE_mouse_genes_WT_infected_vs_WT_not_infected_wilcox_p0.05.xlsx

Additional file 7: DE_mouse_genes_GsdmD_vs_WT_not_infected_wilcox_p0.05.xlsx

Additional file 8: DE_mouse_genes_GsdmD_vs_WT_infected_wilcox_p0.05.xlsx

Additional file 9: DE_mouse_genes_Nlrc4_vs_WT_not_infected_wilcox_p0.05.xlsx

Additional file 10: DE_mouse_genes_Nlrc4_vs_WT_infected_wilcox_p0.05.xlsx

Additional file 11: DEA_Top_20_Molecular_Functions_for_Downregulated_Condition_Nlrc4_WT_infected_BA_organoid_DEGs.pdf

Additional file 12: DEA_top_20_GO_Biological_Processes_for_Upregulated_Condition_WT_infected_BA_organoid_DEG s.pdf

Additional file 13: Colocalisation_wilcox_mouse_BA_filt_norm_de_genes_WT_infected_p0.05.xlsx

Additional file 14: Colocalisation_wilcox_mouse_BA_filt_norm_de_genes_Nlrc4_p0.05.xlsx

Additional file 15: Colocalisation_wilcox_mouse_BA_filt_norm_de_genes_GsdmD_p0.05.xlsx

