## Additional file 3 for "ProbeST: a custom probe design pipeline for Spatial Transcriptomics"

the final amount of probes is:  51 

the total amount of sequences designed probes for:  19
the amount of sequences containing  0  probes:  4
the amount of sequences containing  1  probes:  1
the amount of sequences containing  2  probes:  4
the amount of sequences containing  3  probes:  14
the amount of sequences containing  4  probes:  0
the amount of sequences containing  5  probes:  0
the amount of sequences containing  6  probes:  0
the amount of sequences containing  7  probes:  0
the amount of sequences containing  8  probes:  0
the amount of sequences containing  9  probes:  0
the amount of sequences containing  10  probes:  0
the amount of sequences containing  11  probes:  0
the amount of sequences containing  12  probes:  0
the amount of sequences containing  13  probes:  0
the amount of sequences containing  14  probes:  0
the amount of sequences containing  15  probes:  0
the amount of sequences containing  16  probes:  0
the amount of sequences containing  17  probes:  0
the amount of sequences containing  18  probes:  0
the amount of sequences containing  19  probes:  0
the amount of sequences containing  20  probes:  0
the amount of sequences containing  21  probes:  0
the amount of sequences containing  22  probes:  0
the amount of sequences containing  23  probes:  0
the amount of sequences containing  24  probes:  0
the amount of sequences containing  25  probes:  0
the amount of sequences containing  26  probes:  0
the amount of sequences containing  27  probes:  0
the amount of sequences containing  28  probes:  0
the amount of sequences containing  29  probes:  0
