## Supplementary figures and images for "ProbeST: a custom probe design pipeline for Spatial Transcriptomics"

### Additional file 11

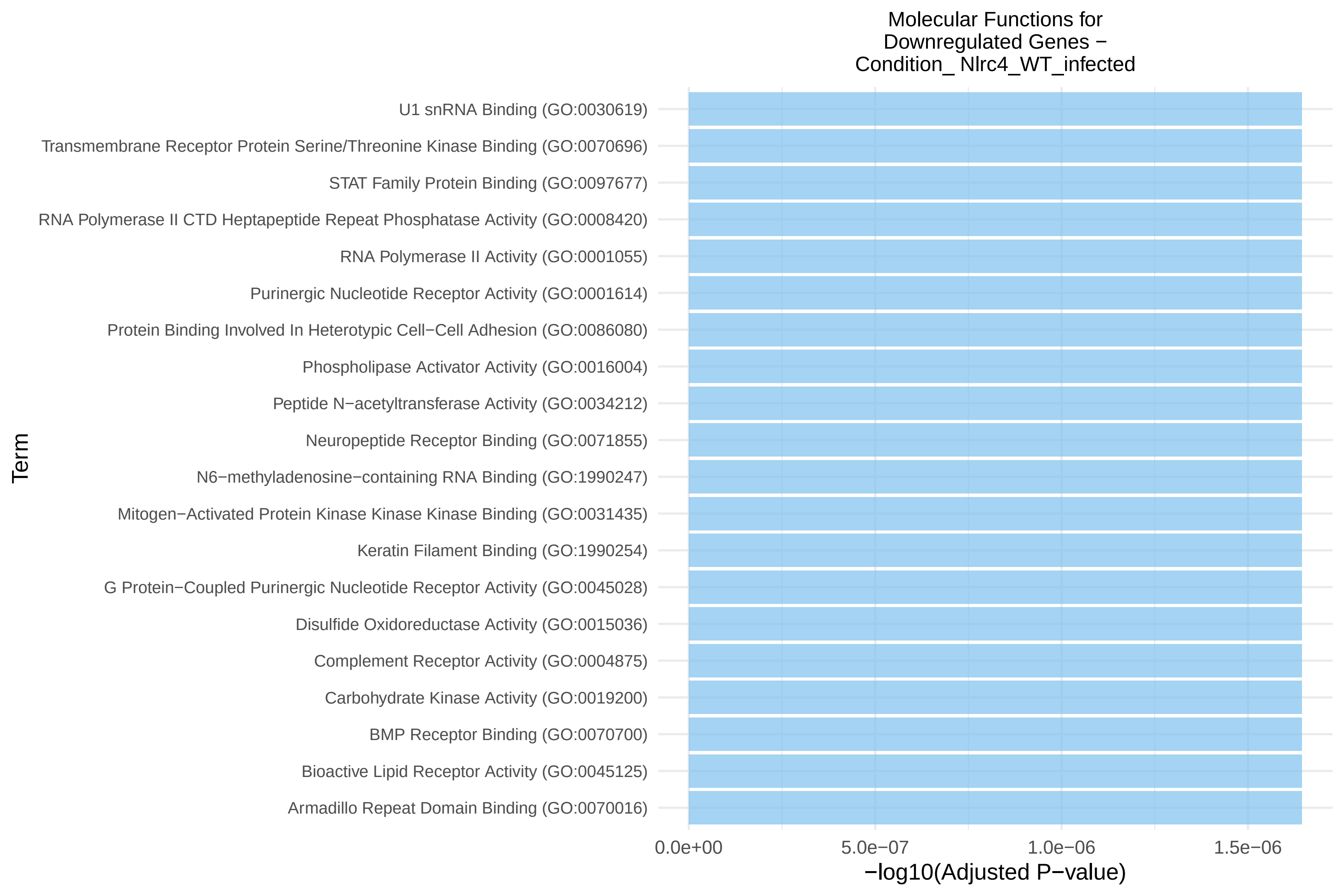

### Additional file 12

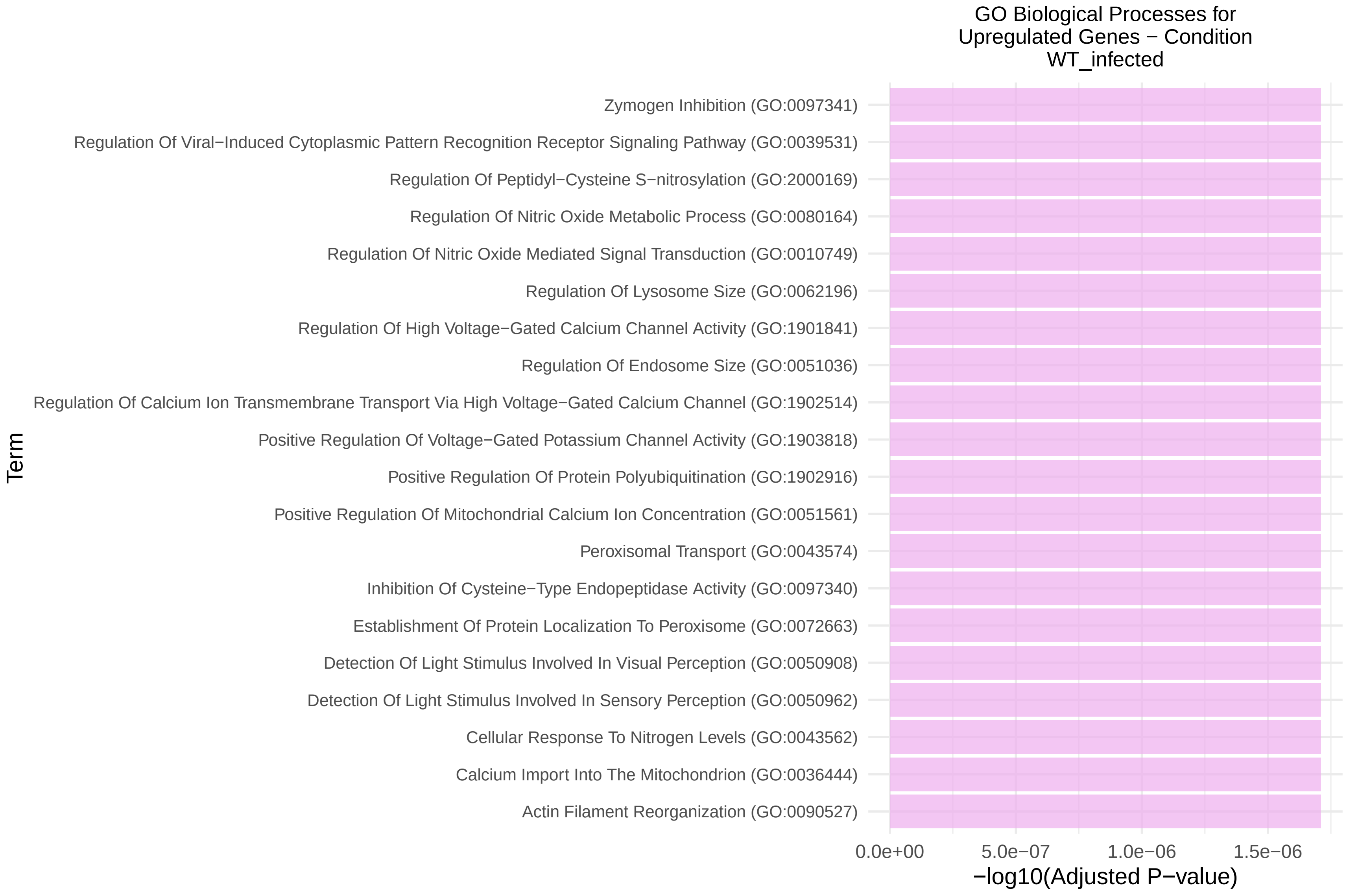
